# Intensity mismatch asymmetry in tinnitus – in which direction should participants pay attention?

**DOI:** 10.1101/2024.01.02.573912

**Authors:** Ekaterina A Yukhnovich, Kai Alter, William Sedley

**Affiliations:** Newcastle University

## Abstract

The effects attention has on intensity deviant Mismatch Negativity responses is an unknown factor in basic sensory neuroscience. It would be useful to understand how attention would affect responses to intensity deviants compared to each other (upward vs downward), and compared to other sensory dimensions such as frequency. Overall, previous research indicates that attention may modulate neuronal gain in healthy participants and change the amplitudes of evoked responses, and may mainly affect the responses to regularly repeating (standard) stimuli rather than deviants. Gain may respond differently in participants with tinnitus and/or hyperacusis under the same conditions compared to controls. Overall, results of the passive task condition were consistent with previous research. Auditory attention magnified MMN in response to upward deviants, while visual attention attenuated it in both control and tinnitus groups. However, auditory attention selectively enhanced downward deviant MMN in the tinnitus group (compared to passive attention). Using the auditory attention paradigm may be advantageous in MMN studies on tinnitus/hyperacusis because the observed differences would be particularly large.

## Introduction

The task performed while a participant is presented with the auditory stimuli during electroencephalography (EEG) may change evoked responses potential (ERP) amplitudes, or even the pattern of ERP responses across different conditions [1]. It is important to understand the effects of attention for a number of reasons, including helping to establish what role attention played in our previous findings, and whether attention is a candidate explanation for why tinnitus and hyperacusis were associated with certain differences in evoked responses profiles. Additionally, it would be useful to establish the optimal attentional state for studying differences due to tinnitus and/or hyperacusis. Finally, the effects attention has on intensity deviant responses is an unknown factor in basic sensory neuroscience. It would be useful to understand how attention would affect responses to intensity deviants compared to each other (upward vs downwards), and compared to other sensory dimensions such as frequency.

Based on Bayesian models of perception, attention may enhance the precision of sensory input [2, 3]. Attentional gain modulation works through top-down modulation of sensory precision, which relies on postsynaptic gain, inhibitory interneurons and feedback mechanisms, the strength of which was positively correlated with attention [3]. Attention may be integral in the formation of optimal predictions in change detection paradigms, but also have dissociable effects from simply strengthening or weakening predictions or prediction errors [2, 4]. For example, an fMRI study used a grating orientation identification task, where a cue determined towards which side of a screen participants needed to direct their attention and respond if a visual stimulus appeared on the attended side [2]. This was done with the aim of studying the role of attention on prediction in the visual cortex. The researchers found that in the unattended condition, standard stimuli showed lower activity than deviants, however, the standards elicited larger responses than deviants when stimuli were attended and task-relevant [2, 5]. In a different study, eighteen healthy participants were tested using an auditory duration deviant oddball paradigm [1], which was presented three times, where participants were asked to either ignore auditory stimuli, passively listen, or focus on the stimuli. Participants were overall younger than those expected to age-match with the tinnitus group in the current study. Additionally, all participants had normal hearing thresholds. The tasks in which participants focused on the stimuli or passively listened showed largest MMN amplitudes, while the smallest amplitudes were in the task where they ignored the stimuli. An important conclusion was made in a review that indicated that direction of attention was more likely to modulate the MMN responses to standard tones rather than the deviants, as MMN reflects the organisation of the prior stimuli in memory as well as the response to the deviant itself [5]. This has also been noticed in some of the previous findings of the current author, where the main differences between conditions was the P200 response to standards but raw deviant forms did not significantly change [6].

Overall, previous research indicates that attention may modulate neuronal gain in healthy participants and change the amplitudes of ERPs, but any effects of attention are typically represented primarily within the modality of the changing stimuli [7], and mainly affect the responses to standard stimuli rather than deviants themselves. It has been tested how attention affects MMN more generally, but not specifically in terms to intensity paradigms. Gain may respond differently in participants with tinnitus and/or hyperacusis under the same conditions, particularly at the tinnitus frequency [8]. In the current study, the auditory system is investigated to see whether there are similarities in the way that attention affects neural prediction error correlates such as MMN, and to see how presence of tinnitus may interact with these attention-related changes. An additional important aim of this study is to understand whether the passive attention condition used in some of the previous MMN-related experiments was adequate [9, 10].

## Materials & methods

### Participants

Volunteers with tinnitus (N=20) were recruited from affiliated volunteer lists at Newcastle University and via Google Ads. General inclusion criteria included being over 18 years old, and able to make an informed choice about volunteering. General exclusion criteria included using ongoing sedating or nerve-acting medications, and mental health conditions severe enough to interfere with everyday life activities. To be included in the tinnitus group, participants had to have chronic tinnitus for over 6 months, with no physical source and that was not due to Meniere’s disease (this was an exclusion criterion for the control group). The control participants were individually matched to participants with tinnitus based on gender, age and an approximate match of their overall audiometric profiles. All participants completed the Hyperacusis Questionnaire [11].

Approval was given by the Newcastle University Research Ethics Committee, and all participants gave written informed consent according to the Declaration of Helsinki (reference number 5619/2020).

### Common methods: tinnitus psychophysics and EEG

The psychophysical assessment in which the tinnitus frequency and the intensity of sounds played during the EEG recording were determined, and the experimental design, followed the procedure in previous EEG experiments within the lab [9, 10]. There were three main differences in the current study. Firstly, in order to reduce the number of factors involved in the analysis, only the tinnitus frequency was included in the paradigm (similarly to the single frequency paradigm in the context study). Secondly, there were gap deviants instead of duration deviants approximately every 1 out of 10 stimuli. These were used as the control deviant condition and as targets in the auditory task. The gap deviants were equally present in every attention block type, the only difference between the tasks was that in the auditory task, participants had to respond to by clicking a keyboard button. Lastly, each participant performed three tasks during their EEG recording, which were presented in randomised order. The tasks were: 1) A 10 minute visual task (x2), in which participants were asked to look at moving dots on the screen and decide whether they are moving randomly or in a specific direction. This task was adapted from code previously created for the Random Dot Kinematograms test [12], with larger gaps between each trial for similarity in task difficulty; 2) A 10 minute auditory task (x2), in which participants were asked to listen out for particularly long gaps (gap deviants within the paradigm) in the stimuli and press a button when they could hear the gap; 3) A 10 minute period (x2) in which participants were asked to watch a subtitled movie of their choice. Independent Component Analysis was used to remove ocular artefacts rather than Denoising Source Separation.

### Statistical Analysis

Statistical analysis was performed using MATLAB. To compare the evoked responses in participants with and without tinnitus, a three-way ANOVA was used with group, intensity, and task as factors of interest, and including interaction terms. Post-hoc analysis included Tukey Honest Significance Tests to determine any significant differences between the ERP amplitudes of the two groups in each task.

## Results

### Demographic information

Table 1 shows means and standard errors (SE) of the demographic information of the two groups, their HQ scores, and THI scores in the tinnitus group. As age was not normally in either group, an independent samples Mann-Whitney U test was used to show that age was not significantly different between the two group (p=0.529). However, HQ scores were significantly different between the groups (p=0.004). Therefore, interpretation of results will have to account for presence of hyperacusis; the score in the tinnitus group is just above the updated cut-off score, but 12 of the 20 participants within the sample scored above the cut-off. Chi-square test was carried out to establish that there were no significant differences in gender across the two groups (Chi (1) =2.50, p=0.114). As the pure tone audiometry results were not normally distributed at the majority of frequencies/ears Mann-Whitney U tests were performed and showed that while 0.25 kHz and 0.5 kHz pure audiometry results were significantly different between the two groups, there were no significant differences between groups at 1, 2, 4, 6 or 8 kHz in right or left ears, with the smallest p-value being p=0.277. The discrepancy at the lowest measured frequencies should not be problematic for this study, as tinnitus usually appears at high frequencies and both groups had mean hearing thresholds within normal range at 0.25 and 0.5 kHz in both ears.

**Table 1.** Descriptive statistics of the study groups. Means and standard errors are given for every group for their age, HQ scores and THI scores. The gender split is also indicated for each group.

|  | Age | Gender | HQ Scores | THI Scores |
| --- | --- | --- | --- | --- |
| **Tinnitus** | 56.20 (3.52) | 13 f, 7 m | 16.05 (1.78) | 34.20 (4.33) |
| **Control** | 57.90 (3.60) | 12 f, 8 m | 9.65 (1.34) | N/A |

### Time course of the stimulus response

Grand average ERP data for channel FCz (with P9/P10 reference) across standard and deviant responses for all stimulus conditions and in each group was used to determine timeframes for quantifying P50, N100, P200 and MMN responses, based on visual inspection (Fig 1). To calculate the MMN difference waveform, standard responses were subtracted from their equivalent deviant conditions.

**Fig 1.**
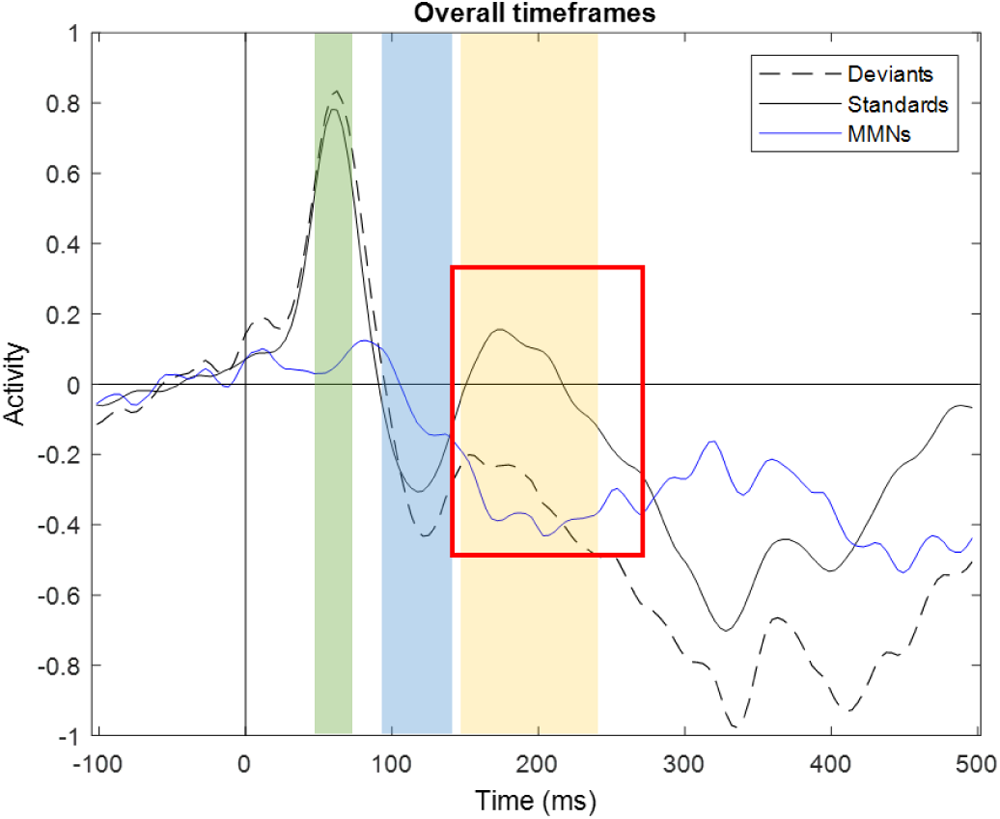
Standard, deviant and MMN waveforms broadly combined across groups and conditions. The timeframe for quantifying P50 was 50-75 ms (green), N100 was 95-140 ms (blue), P200 was 150-240 ms (orange). The MMN timeframe was 140-260 ms (red square). As the MMN and P200 overlap, P200 was only used in standards (and MMN in the difference waveform).

### Standard and pure deviant waveforms

The average waveforms, split by task and intensity, are shown on Fig 2.

**Fig 2.**
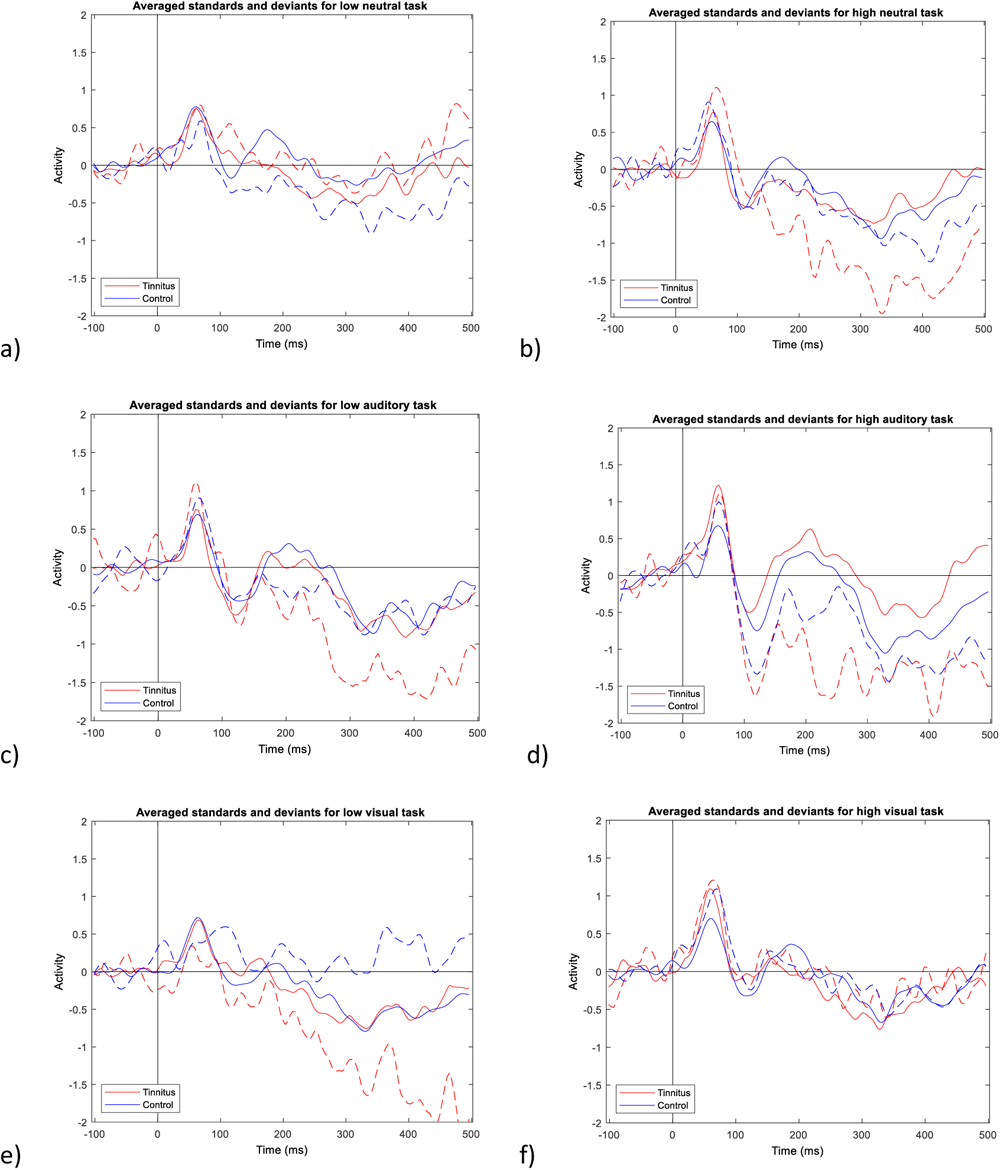
Average waveforms to standard and deviant tones in different intensity and task conditions. The tasks, from top to bottom, are neutral, auditory, and then visual. On the left, are the quieter stimulus conditions and on the right are the louder stimulus conditions. Standard responses are shown in solid lines and deviants are shown in dashed lines. The groups are colour-coded as follows: tinnitus (red), controls (blue).

### P50 responses

Fig 3 shows P50 responses to standard and deviant stimuli in the three tasks at two intensities. A three-way ANOVA (group, task, intensity) was carried out on P50 responses to standard tones, which found main effects of group and intensity (both p<0.001), but not task (p=0.091). There were significant interactions between group and intensity, and task and intensity (both p<0.001). There were also significant interactions between group and task (p=0.005) and group, task and intensity (p=0.031). From visual inspection (in keeping with the reported statistical effects), the biggest difference was that the two active tasks increased P50 to high intensity stimuli in the tinnitus group, reflecting a combination of these interaction effects. Post hoc tests showed that there were no significant differences between P50 response amplitudes to the three tasks at the low intensity. However, the neutral task had significantly smaller amplitudes than the other two tasks (p<0.001), driven by the tinnitus group having larger responses than the controls (p=0.011 for auditory and p<0.001 for visual).

**Fig 3.**
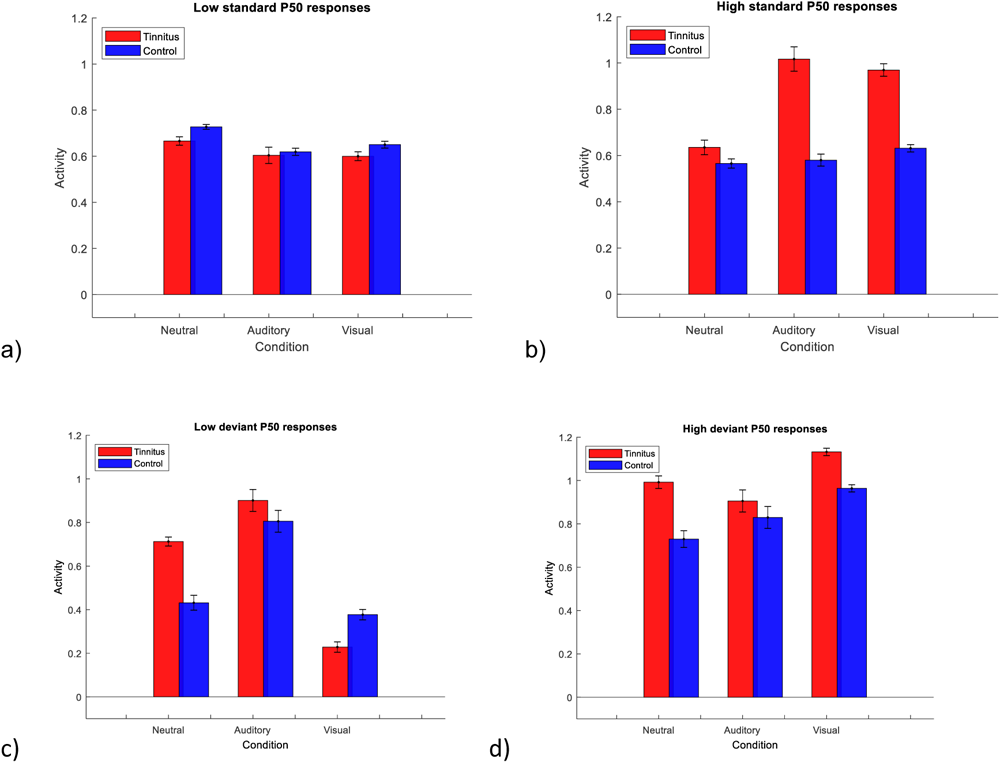
Standard and deviant responses in the P50 timeframe. Top two charts represent averaged responses to standard stimuli and bottom charts represent averaged responses to deviant stimuli. Tinnitus group is represented in red and controls are represented in blue.

When comparing the pure deviant responses, a three-way ANOVA (group, task, intensity) showed main effects of group, task and intensity, and a significant interaction of task and intensity (all p<0.001). In response to the low deviant during the auditory task, both groups had larger P50 amplitudes than during either of the other tasks. Additionally, during the visual task, both groups had particularly lower P50 amplitudes in response to low deviant compared to high deviant.

There was also a significant interaction effect between group and task (p=0.0.004). Tinnitus group had significantly higher P50 amplitudes compared to controls in response to low deviant in neutral task (p=0.002) and significantly lower P50 amplitudes in response to low deviant in visual task (p=0.004). Based on visual observation of Fig 2, the visual low deviant likely showed controls to have larger P50 amplitudes than the tinnitus group because there was no reduction of the amplitude due to lack of N100 in controls but not in the tinnitus group.

### N100 responses

Fig 4 shows the mean amplitudes of N100 responses to standard and deviant tones in the three tasks. A three-way ANOVA (group, task, intensity) showed main effects of all three factors, and interactions between task and intensity, and group, task and intensity (all p<0.001). There was also a significant interaction between group and task (p=0.022). At the low intensity standards, N100 amplitudes for the auditory task were significantly lower than for both neutral and visual task (p<0.001 and p=0.006, respectively). At high intensity standard, N100 amplitudes were significantly weaker in the visual task than neutral and auditory tasks (p<0.001). Post hoc tests showed significantly weaker responses in tinnitus group compared to controls within the neutral and visual tasks at low intensity standards (p=0.003 and p=0.002, respectively), and at high intensity standards within auditory and visual tasks (p<0.001 for both). Based on visual inspection of Fig 2, the tinnitus group did not have a clear N100 in response to low standards in the neutral task. The tinnitus group had non-significantly stronger N100 amplitudes than the controls for the low standard in the auditory task and for the high standard in the neutral task.

**Fig 4.**
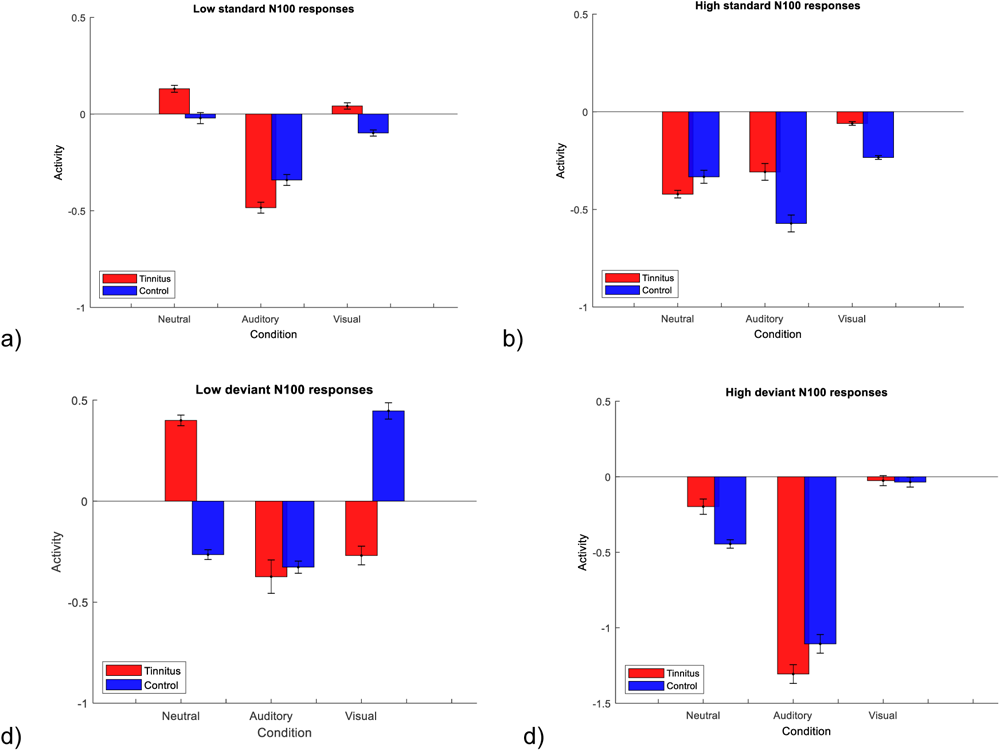
Standard and deviant responses in the N100 timeframe. the top two charts represent averaged responses to standard stimuli, bottom charts represent averaged responses to deviant stimuli. The order of the groups in all charts is: neutral task, auditory task, visual task.

For deviant tones, a three-way ANOVA (group, task, intensity) showed significant main effects of task and intensity, as well as significant interactions between group and task, task and intensity, and group, task and intensity (all p<0.001). At the low intensity deviant, the tinnitus group showed significantly weaker N100 than the controls in the neutral task (p<0.001) but significantly stronger N100 than controls in the visual task (p<0.001). For the high deviant, the auditory task elicited significantly stronger responses than the other two tasks overall (p<0.001), with the visual task showing the lowest amplitude for N100 (p<0.001 compared to the others). In the neutral task, the tinnitus group had significantly lower N100 amplitude than the controls (p<0.001).

### P200 responses

Fig 5 shows the mean amplitudes of P200 responses to standard tones in the three tasks. A three-way ANOVA (group, task, intensity) showed main effects of all three factors (p=0.036 for group, p<0.001 for task and intensity), as well as an interaction between group and task, task and intensity, group task and intensity (all p<0.001), and group and intensity (p=0.009). In response to both intensities during the neutral task, the tinnitus group had significantly lower amplitudes than the controls (both p<0.001). For high intensity standards during the auditory task, there was a significantly larger response in the tinnitus group compared to controls (p<0.001), possibly due to the prior N100 amplitudes. Additionally, the controls had a significantly stronger response to high standards in the visual task (p<0.001). At both intensities, the tinnitus group had higher P200 responses to the auditory task than neutral or visual tasks (p<0.001), and lower P200 amplitude in the high intensity responses during the neutral task (p<0.001).

**Fig 5.**
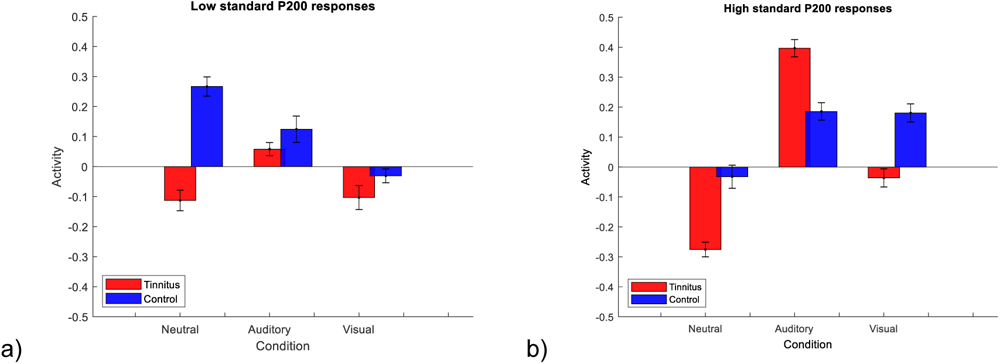
Standard responses in the P200 timeframe. The order of the groups in all charts is: neutral task, auditory task, visual task.

### Mismatch Negativity (MMN)

A difference waveform was calculated by subtracting standard waveforms from deviant waveforms, which was used for MMN analysis (Fig 6).

**Fig 6.**
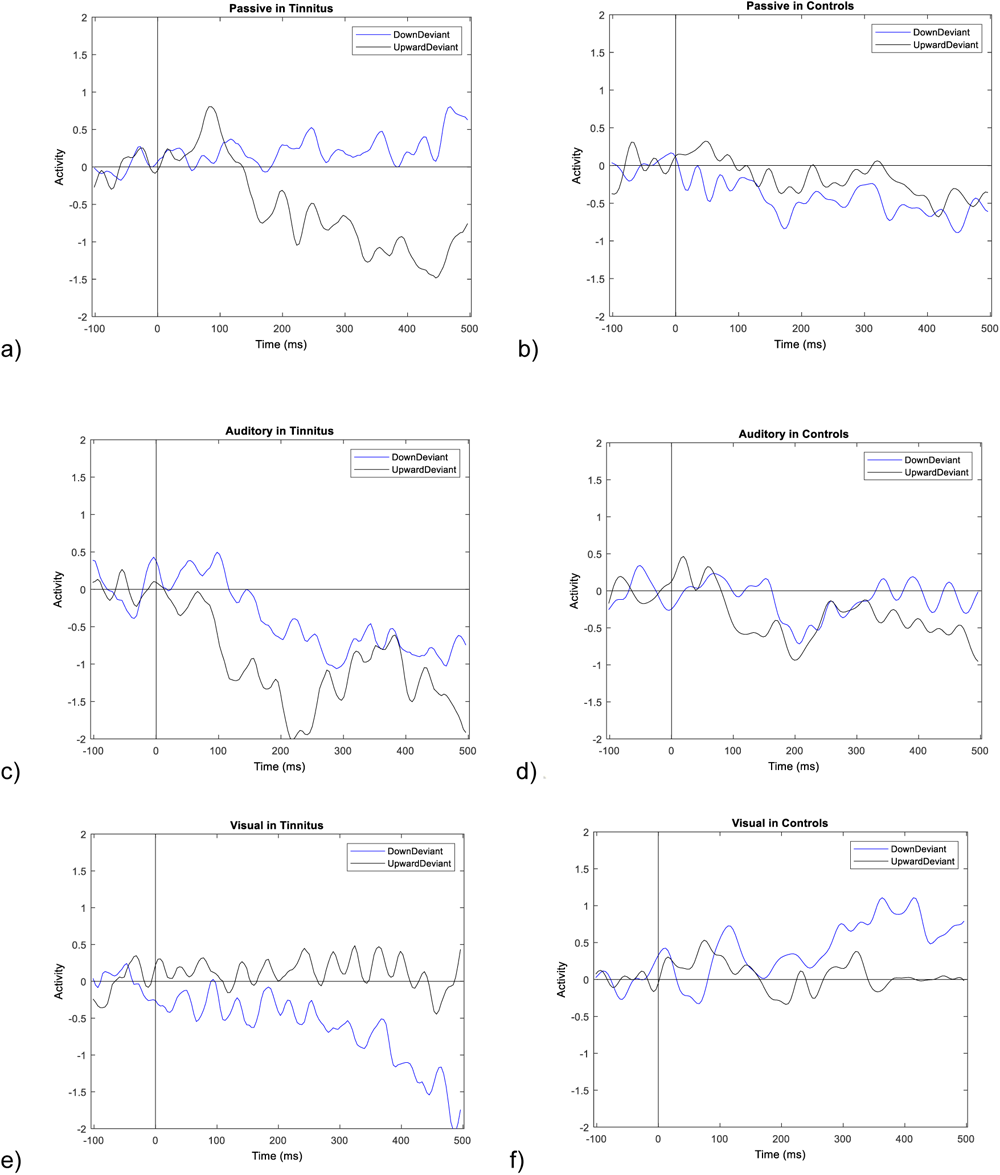
Average difference waveforms in different intensity and task conditions. The tasks, from top to bottom, are neutral, auditory, and then visual. The tinnitus group is on the left and the controls are on the right. The conditions are colour-coded as follows: DD (blue), UD (black).

Fig 7 shows MMNs in the three tasks. A three-way ANOVA (group, task, intensity) showed significant main effects of all three factors, as well as significant interactions between all pairs and between all three factors (all p<0.001). Post hoc tests showed that at each intensity and in each task, except DD for the auditory task (p=0.058), there was a significant difference between tinnitus and control groups (all p<0.001), though the largest difference was in the auditory group in response to UDs. Tinnitus group had weaker MMNs compared to controls only for DDs but stronger MMNs for UDs in the neutral task, but the opposite pattern was seen for MMNs during the visual task.

**Fig 7.**
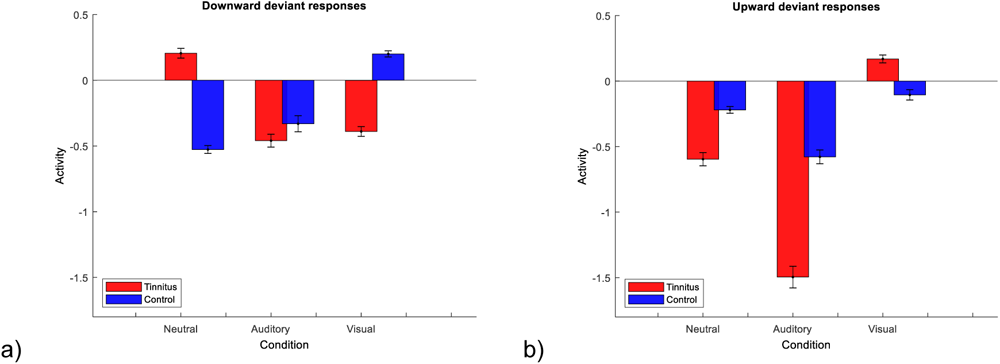
MMN responses. The order of the groups in all charts is: neutral task, auditory task, visual task.

## Discussion

### Main findings

Results of the passive task condition were consistent with previous studies. Auditory attention magnified MMN in response to upward deviants, while visual attention attenuated it. Auditory attention selectively enhanced downward deviant MMN in the tinnitus group (compared to passive attention).

### P50s

P50 responses were similar in response to low standards in both groups across the three tasks. While the responses remained similar between the groups for the high standard in the neutral task, in the visual and auditory tasks the tinnitus group had larger P50 amplitudes, while control group responses remained similar to each other and to low standard. The differences in response amplitude in the tinnitus group was probably not due to the presence of hyperacusis because, as shown in the findings in [6], T+H+ group had weaker P50 amplitudes both near and outside of tinnitus frequency (though there is also a possibility that results in the two studies may not be comparable due to the overall context of the paradigm). Regarding the overall context, it is also possible that the tinnitus group in the current study had less overall/less quick suppression to the repeating sound when they had to either focus on the stimuli, or the stimuli were interfering with focus on a task in a different modality. Participants with tinnitus may have had more difficulty gating out the stimuli compared to the control group during the attention-demanding tasks due to less ability to ignore the irrelevant stimuli [13, 14], but not in the neutral task where they did not need to pay attention to the stimuli.

Controls had larger high deviant responses and smaller low deviant responses compared to standards, except during the auditory task where low deviant responses were similarly increased to high deviant responses. Tinnitus group showed similar patterns to the controls, but neutral condition responses appeared more like auditory condition in response to low deviant. Additionally, the tinnitus group also had larger deviant P50s to both intensity directions during the neutral task. So, while tinnitus participants were not concentrating on the stimuli or on a particular task, they were more responsive to changes in the stimuli. This was further shown by the tinnitus group having higher P50s for all tasks in response to high deviants. In the auditory task, P50 amplitudes were similar for both groups in response to both intensity deviants.

As it has been suggested that P50 amplitudes reduce for repetition of the immediately preceding tone, it is possible that the tinnitus group may have reduced inhibitory function compared to controls when attention is manipulated [15, 16]. Further, while much of previous literature showed that P50 was not dependent on attention (unlike N100), P50 amplitudes may be altered by psychological stressors, both in terms of task (e.g. mental arithmetic) and more chronic difficulties such as PTSD [17, 18]. So, perhaps either hyperacusis presence or tinnitus itself relates to higher stress responses both when paying attention or purposefully ignoring tinnitus-like stimuli.

The control group responses in the neutral task were similar to results seen in response to the small frequency difference paradigm in [19].

### N100s

The tinnitus group showed a similar (almost) fully suppressed N100 in response to standard tones during the low intensity condition in the neutral task and during both intensity conditions in the visual task. In the neutral task, suppression was only present during the low standard for tinnitus, which could occur because the higher intensity stimuli drew more attention in a stimulus-driven manner in a way that does not occur with lower or decreasing stimuli. N100 has been regarded as an attention-triggering mechanism [20]. This suppression may not have been present in the auditory task because participants were focusing on the stimuli; however, the more attention was taken away from the stimuli, the less sensitive the neurons became (less gain) and the more suppression was present [2], and this did not occur in the controls because they were not used to having to ignore an auditory stimulus while completing tasks. As possible corroboration of this explanation, a previous MEG study showed that actively ignoring a particular auditory stimulus was related to reduced M100 amplitudes compared to passive condition [21].

In the auditory task, the tinnitus group had stronger responses to the low standard but controls had stronger responses to high standard stimuli. At the high standard (but not low), this could have occurred due to the prior P50 amplitudes, which in turn may have occurred due to hyperacusis, and subsequently affected P200 in the standards. However, N100 and P200 are not always dependent on each other, again pointing towards potentially this paradigm being more useful for sensory gating/ adaptation experiments into tinnitus and/or hyperacusis rather than MMN [22].

N100 amplitudes were present and stronger overall, particularly in response to high deviants in the auditory task compared to the other two tasks, likely due to the concentration on the stimuli. In response to low deviants, N100 was not clear for the tinnitus group during the neutral task (but it was present in controls); in high deviants, N100 was also weaker in tinnitus group than controls. The opposite pattern occurred in response to low deviants the visual task, where the tinnitus group showed an N100 but controls did not, and both groups had weak responses to the high deviants. It would be interesting to understand why these pattern swaps occur.

### P200s

P200 was more present in controls than in the tinnitus group, except during auditory task in response to high standards. Controls had more positive P200 response to the neutral task in low standards, but to auditory and visual tasks in high standards. In the tinnitus group, P200 was more positive during the auditory task overall, and particularly in response to high standards. P200 has been associated with allocation of attention [22]. Previous studies have found reductions in P200 amplitudes in tinnitus participants compared to controls [23], though other studies saw differences in latencies but not amplitudes [24]. P200 amplitude could be manipulated through task demands, as well as stimulus parameters [25]. P200 tends to be larger when attentional demands are lower [26]. This may explain why P200 was larger for tinnitus group during the auditory task compared to the other tasks in both intensity conditions: they only needed to pay attention to the auditory stimuli, which already takes some of their attention as part of tinnitus. However, both neutral and visual tasks required some focus on other modalities and tasks. Control responses, however, are harder to explain using this theory. Differences in N100 may be obscuring the relative relationships between high and low intensities.

### MMNs

In the neutral task, the tinnitus group had larger responses to the upward deviants and smaller responses to downward deviants, while controls had the opposite pattern of MMN amplitudes. This finding is similar to the original study [10] and the T+H+ group in [6].

In the auditory task, controls did not show much difference from the neutral task, but there was a slight increase in response to upward deviants compared to downward deviants. This difference was seemingly mediated by differences in standard P200 responses [5]. In the tinnitus group, larger responses were seen to downward deviants compared to controls. This was not driven by P200 standard responses, and perhaps was in keeping with the result of T+H-group in [6], where increased MMN to downward deviants was the sole MMN change related to tinnitus once hyperacusis had been controlled for. The larger response to the upward deviant may have been driven by P200. Aside from P200 findings, the main results were that the tinnitus group had larger downward deviant MMN in the auditory task than the passive condition.

In the visual task, MMN responses to upward deviants were largely attenuated. In response to downward deviants, MMN was a somewhat more present for the tinnitus group compared to upward deviant, while for control group MMN was abolished even compared to upward deviant.

The oddball paradigm in a previous MMN attention study mentioned in the introduction was similar to the current study [1]. The tasks in which participants focused on the stimuli or passively listened showed the largest MMN amplitudes, while the smallest amplitudes were in the task where they ignored the stimuli. A difference between the previous and the current studies was that previously, all paradigms elicited significant MMNs while this was not the case in the current study. It is possible that duration deviants elicited somewhat stronger responses than the intensity changes in the current study, or even that the overall context of their study included word and sentence paradigms within the same session, all of which were used to elicit MMNs, and that may have changed the waveform shapes somewhat. A potential explanatory factor in the diminished responses, particularly to the visual task, could be stress or arousal, as this task likely involved higher cognitive load due to involvement of different modalities, For example, in a previous study, cortisol level was inversely related to MMN amplitudes in response to duration deviants, and psychosocial stressors overall attenuated processing of change detection [27].

There was an inverse relationship between deviant P50s and MMN in all groups in response to all conditions [28]. This was particularly present in responses to the upward deviants, where during the visual task, P50s were the highest compared to the other two task, and MMNs were abolished. This relationship was still present in response to downward deviant, but it was somewhat less apparent, possibly due to the overall lower amplitude in response to the quieter tones. However, it may be interesting to further investigate this relationship.

### Paradigm recommendations

Overall, auditory attention increased P50 responses to downward deviants, facilitated N100 formation to low intensity standards, facilitated P200 formation overall, increased upward deviant MMNs as an effect of standard P200 rather than deviant waveform, and increased N100 and MMN responses to upward deviants. In tinnitus, auditory attention exaggerated P200 formation, exaggerated MMN enhancement to upward intensity deviants, and enhanced MMN to downward intensity deviants. It was shown that when paying attention to auditory stimuli, participants with tinnitus (though more likely this was due to hyperacusis) were particularly more responsive than controls to a change in the stimulus in the upward intensity direction. This finding was similar to the T+H+ group in [6]. It may be useful to investigate whether there would be stronger difference between the groups in response to the downward deviant condition in a tinnitus sample without hyperacusis.

Using the auditory attention paradigm may be advantageous in MMN studies on tinnitus/hyperacusis because the observed differences would be particularly large. Similarly, this task would be useful in investigating sensory gating or habituation processes due to its effect on P50, N100 and P200 amplitudes. However, to avoid both boredom/fatigue and extreme repetition positivity, the length of the experiment would need to be shortened, or some breaks would need to be implemented.

Passive listening (in the neutral task) gave overall similar response profiles, though often much weaker, to auditory attention, meaning that one can generally compare the results of attended and passive studies meaningfully. Therefore, results of the previous IMA studies [9, 29], in which passive attention task was utilised, were unlikely to be due to attentional differences in tinnitus and/or hyperacusis, because attention to auditory stimuli in the present study exaggerated differences between tinnitus and control groups rather than causing or abolishing them. The neutral task elicited most similar standards across the two groups at P50 and N100 ERPs, while the auditory condition showed the strongest responses on average. Therefore, responses to neutral task would probably be the most representative for later components such as P200 or MMN, however they will not be as strong as in the auditory attention task. For particularly long paradigms, or those that cannot include breaks, this may be the more optimal paradigm, as it would show similar, albeit less striking results compared to the auditory paradigm and would not be as potentially taxing for participants.

Visual attention largely had the opposite effects to auditory attention: abolishing N100, especially to high intensity standards and deviants, and MMN. However, in some limited instances in tinnitus subjects it had similar effects to auditory attention (increasing P50 to high standard stimuli, and increasing N100 and MMN to downward deviants), and these changes may be more indicative of state of arousal/stress/readiness/task-engagement than sensory attention per-se. As such, this paradigm may be used to study the distress-related symptoms or responsiveness to stimuli in tinnitus or possibly hyperacusis.

Overall, the results of this study could occur due to alterations of central gain, such as enhancement with directed attention and inhibition with an interfering task [2]. Within the Free Energy formulation, attentional gain modulation works through top-down modulation of sensory precision, which is controlled by post-synaptic gain [3]. Attention acts by increasing post-synaptic gain/precision, to the point where they can be considered interchangeable. There might have been some differences in adaptation mechanisms between groups, including altered dynamic range adaptation and sensory gating in the tinnitus participants, which was affected both by task and intensity conditions. Dynamic range adaptation of neural firing controls the excitability of the neurons to a desired rate, and therefore improving efficiency of neural coding of intensity (both increasing and suppressing activity in response to mean sound level in the environment) [29]. A reduced dynamic range leads to increased sensitivity to sound intensity, which has been found in people with tinnitus/hyperacusis [30]. While participants could adjust the loudness of the sounds they listened to, they may have continued to be more responsive to the presence of the sounds, as indicated through less sensory gating. In that case, the comparisons made by the error units in participants with tinnitus would not follow the patterns of controls. The results seen in this study could be due to the different weighting participants with tinnitus assign to prediction errors, as well as different functioning of sensory gain and inhibitory interneurons. As such, in the future researchers should be intentional with their choice of modulation of attention during depending on their primary focus.

